# Sensitive and reproducible cell-free methylome quantification with synthetic spike-in controls

**DOI:** 10.1101/2021.02.12.430289

**Authors:** Samantha L. Wilson, Shu Yi Shen, Lauren Harmon, Justin M. Burgener, Tim Triche, Scott V. Bratman, Daniel D. De Carvalho, Michael M. Hoffman

**Author notes:** Authors contributed equally to this work (Samantha L. Wilson, Shu Yi Shen).

## Abstract

**Background:** Cell-free methylated DNA immunoprecipitation-sequencing (cfMeDIP-seq) identifies genomic regions with DNA methylation, using a protocol adapted to work with low-input DNA samples and with cell-free DNA (cfDNA). This method allows for DNA methylation profiling of circulating tumour DNA in cancer patients’ blood samples. Such epigenetic profiling of circulating tumour DNA provides information about in which tissues tumour DNA originates, a key requirement of any test for early cancer detection. In addition, DNA methylation signatures provide prognostic information and can detect relapse. For robust quantitative comparisons between samples, immunoprecipitation enrichment methods like cfMeDIP-seq require normalization against common reference controls.

**Methods:** To provide a simple and inexpensive reference for quantitative normalization, we developed a set of synthetic spike-in DNA controls for cfMeDIP-seq. These controls account for technical variation in enrichment efficiency due to biophysical properties of DNA fragments. Specifically, we designed 54 DNA fragments with combinations of methylation status (methylated and unmethylated), fragment length (80 bp, 160 bp, 320 bp), G+C content (35%, 50%, 65%), and fraction of CpG dinucleotides within the fragment (1/80 bp, 1/40 bp, 1/20 bp). We ensured that the spike-in synthetic DNA sequences do not align to the human genome. We integrated unique molecular indices (UMIs) into cfMeDIP-seq to control for differential amplification after enrichment. To assess enrichment bias according to distinct biophysical properties, we conducted cfMeDIP-seq solely on spike-in DNA fragments. To optimize the amount of spike-in DNA required, we added varying quantities of spike-in control DNA to sheared HCT116 colon cancer genomic DNA prior to cfMeDIP-seq. To assess batch effects, three separate labs conducted cfMeDIP-seq on peripheral blood plasma samples from acute myeloid leukemia (AML) patients.

**Results:** We show that cfMeDIP-seq enriches for highly methylated regions, capturing ≥ 97% of methylated spike-in control fragments with ≤ 3% non-specific binding and preference for both high G+C content fragments and fragments with more CpGs. The use of 0.01 ng of spike-in control DNA in each sample provided sufficient sequencing reads to adjust for variance due to fragment length, G+C content, and CpG fraction. Using the known amount of each spiked-in fragment, we created a generalized linear model that absolutely quantifies molar amount from read counts across the genome, while adjusting for fragment length, G+C content, and CpG fraction. Employing our spike-in controls greatly mitigates batch effects, reducing batch-associated variance to ≤ 1% of the total variance within the data.

**Discussion:** Incorporation of spike-in controls enables absolute quantification of methylated cfDNA generated from methylated DNA immunoprecipitation-sequencing (MeDIP-seq) experiments. It mitigates batch effects and corrects for biases in enrichment due to known biophysical properties of DNA fragments and other technical biases. We created an R package, `spiky`, to convert read counts to picomoles of DNA fragments, while adjusting for fragment properties that affect enrichment. The `spiky` package is available on Bioconductor (https://bioconductor.org/packages/spiky) and GitHub (https://github.com/trichelab/spiky).

## 1 Introduction

Cell-free methylated DNA immunoprecipitation-sequencing (cfMeDIP-seq) identifies DNA methylation using low-input samples of cell-free DNA (cfDNA). This method detects DNA methylation patterns reflective of distinct cancer types from circulating tumour DNA, which arises from tumour cells shedding DNA into an individual’s blood.^1,2^ cfMeDIP-seq proves ideal when assessing peripheral blood plasma from cancer patients, where one may obtain only a small amount of circulating tumour DNA and no indication of from where the circulating tumour DNA originates.

Sequencing assay methods, such as cfMeDIP-seq or RNA-seq, require a control for more accurate quantitative comparison across samples and batches. Reference controls for sequencing assays have consisted of spike-in reference DNA or RNA fragments of known sequence.^3–7^ Without spike-in controls, one must assume that a given amount of assayed material produces equal DNA or RNA yields in different experimental conditions, and that this also holds true across all genomic regions.^4^ By normalizing quantification to a known amount of spike-in DNA added into a sample, we can overcome this assumption, leading to more accurate results.^4^ When carefully designed, spike-in controls can adjust for specific technical biases.

In addition to adjusting for technical biases, spike-in controls act as experimental standards for quality control. The addition of spike-in controls dramatically changes the interpretation of RNA-seq, chromatin immunoprecipitation-sequencing (ChIP-seq), and other genomic assay results.^3–7^ As such, all quantitative genome-wide assays would benefit from the addition of spike-in controls.^4^

The most common approach to normalizing sequencing assay data consists in dividing the number of reads at each genomic region by the total number of reads genome-wide. This approach addresses technical variance due to sequencing depth, but it can mask differences in biological variables of interest. Normalizing data to a known amount of spike-in DNA for each sample allows for more accurate detection of differences and adjustment of biophysical properties of DNA fragments that can influence results.^4,5^

While other genomic assays have long utilized spike-in controls, methods measuring genome-wide DNA methylation have rarely used them. The previous cfMeDIP-seq protocol uses methylated and unmethylated *Arabidopsis thaliana* DNA as a spike-in control to assess immunoprecipitation and binding efficiency to methylated DNA.^2^ This approach cannot correct for properties of specific methylated DNA fragments likely to influence results, such as G+C content, fragment length, and CpG fraction. Other controls used in DNA methylation enrichment methods include setting aside and sequencing a portion of input DNA without the enrichment procedure.^8^ This provides a reference point to assess enrichment of DNA methylation overall. Sequencing input DNA, however, cannot adjust for properties of individual fragments that can affect enrichment. Spike-in controls for bisulfite conversion methods for assaying DNA methylation^9,10^ have no use in bisulfite-free enrichment methods, like cfMeDIP.

Here, we introduce new synthetic DNA spike-in controls for cfMeDIP-seq. Our synthetic spike-ins can measure how DNA fragment properties, such as length, G+C content, and number of CpGs, can affect the number of reads produced by some known amount of fragment. Our spike-in controls assess non-specific binding, an integral part of cfMeDIP-seq analysis. The spike-in controls also mitigate technical effects such as experiments performed by different labs. We also use unique molecular indices (UMIs) to adjust for polymerase chain reaction (PCR) bias. To calculate methylation specificity, we compare methylated fragments to unmethylated fragments after cfMeDIP-seq. We add a known molar amount of spike-in controls to each sample. With this information, we apply a generalized linear model to calculate molar amount, accounting for fragment length, G+C content, and CpG fraction. These spike-in controls generate an absolute quantitative measure of methylated DNA, allowing for more robust comparisons between samples and experiments.

## 2 Methods

### 2.1 Designing synthetic DNA spike-in controls

We used public paired-end, whole genome cfDNA sequence data to assess typical cfDNA fragment properties including fragment length, G+C content, and the number of CpG dinucleotides.^11^ We considered the number of CpGs as a fraction of fragment length.

From the observed distribution of cfDNA fragment properties, we set the following spike-in fragment parameters:

- 3 fragment lengths: 80 bp, 160 bp, and 320 bp
- 3 G+C contents: 35%, 50%, and 65%
- 3 CpG fractions: 1/80 bp, 1/40 bp, and 1/20bp

These parameters generate 27 fragment combinations (3 fragment lengths ×3 G+C contents ×3 CpG fractions = 27).

We set the CpG fraction parameters so that every fragment length would have an integer number of CpGs. For example, the 80 bp fragments have 1, 2, or 4 CpG dinucleotides, and the 160 bp fragments have 2, 4, and 8 CpG dinucleotides. We used GenRGenS version 2.0^12^ to construct 27 different first-order Markov models that generate sequences with the desired parameters. For each Markov model, we generated numerous sequences. We then identified those sequences that fulfilled two criteria: (1) no alignment to human genome and (2) no potential secondary structures.

Using blastn,^13^ we searched for alignment of the generated sequences to the human reference genome (GRCh38/hg38).^14^ We ensured no alignment of each synthetic sequence to the genome, and selected the sequences with the lowest E-values in each search.

We used UNAFold software^15^ (Integrated DNA Technologies (IDT), Coralville, IA, USA) to check for secondary DNA structure for 80 bp and 160 bp fragments. For 320 bp fragments, we used RNAstructure version 6.2^16^ to check for secondary DNA structures.

We attempted to pick two sequences from those fulfilling our criteria for each of the 27 Markov models. This represented one sequence for a methylated fragment, and one for an unmethylated fragment. If possible, this would have produced 54 desired spike-in control sequences. The generated 320 bp fragments with 65% G+C content, however, had excessive amounts of secondary structure, which would make synthesis difficult and might have systematic effects on the cfMeDIP process. To retain a total of 18 generated spike-in control sequences for 320 bp fragments, we designed an additional alternative set of fragments with different sequences for each of 3 combinations of G+C content and CpG fraction: (1) 35% G+C content, 1/80 bp CpG fraction; (2) 50% G+C content, 1/40 bp CpG fraction; and (3) 50% G+C content, 1/20 bp CpG fraction.

### 2.2 Synthetic fragment preparation

We acquired synthetic fragments for the spike-in control sequences from a commercial service (IDT, Coralville, IA, USA). For the 80 bp and 160 bp fragments we used 4 nmol Ultramer DNA Oligonucleotides. For the 320 bp fragments we used gBlocks Gene Fragments, obtaining 250 ng of each designed fragment. The two 160 bp fragments with 35% G+C content and 1/20 bp CpG fraction failed commercial design procedures, leaving us with 52 spike-in control fragments (Supplementary Table 1).

We amplified the synthetic fragments by PCR. We used the High-Fidelity 2X Master Mix (New England Biolabs, Ipswich, MA, USA, Cat #M0492L) and the fragments’ optimal annealing temperatures (Supplementary Table 1). We purified amplified fragments using the QIAquick PCR Purification Kit (Qiagen, Hilden, Germany, Cat #28104). We determined concentration of each synthetic fragment via NanoDrop (Thermo Fisher Scientific, Waltham, MA, USA).

For each fragment designed for methylation, we took 1 μg of synthetic DNA fragment, and methylated using *M*.*SssI* CpG methyltransferase (Thermo Fisher Scientific, Waltham, MA, USA, Cat #EM0821). We incubated the methylation reaction at 37 °C for 30 min, then 65 °C for 20 min. We purified the methylated product using the MinElute PCR Purification Kit (Qiagen, Hilden, Germany, Cat #28004). To verify methylation, we digested the original PCR amplicon and the methylated PCR amplicon with either HpyCH4IV, HpaII or AfeI restriction enzyme, depending on the cut sites each fragment contained (Supplementary Table 1). We considered methylation successful when, after restriction digest, the PCR amplicon had a single band when run on a 2% agarose electrophoresis gel. Then, using the known relative molecular mass of each synthetic fragment, we determined its molar amount using the Qubit dsDNA HS Assay Kit and a Qubit 2.0 Fluorimeter (Thermo Fisher Scientific, Waltham, MA, USA).

### 2.3 Assessing technical bias

To assess the performance and any potential biases of synthetic fragments as spike-in controls, we performed cfMeDIP-seq using solely the spike-in control DNA pools as the input. The input pool consisted of 9.99 ng synthetic spike-in DNA, with equimolar amounts of each fragment size within each fragment size pool, and equimolar amounts of each methylation status (Supplementary Table 1).

We performed cfMeDIP-seq as previously described^2^ with slight modifications, detailed here. To account for PCR amplification bias, we used xGen Stubby Adapter and unique dual indexing (UDI) primer pairs (IDT, Coralville, IA, USA, Cat #10005924). We performed adapter ligation overnight at 4 °C, adjusting final adapter concentration to 0.09 μmol by dilution.

For each sample, we saved 10% of the DNA denaturation product, prior to incubation with the antibody, as input (Figure 1, samples 1A and 2A). For each sample, we amplified both input and outputs followed by purification and dual size selection using AMPure XP beads (Beckman Coulter, Brea, CA, USA) selecting for fragments between 200 bp–500 bp, reflecting the addition of adapter to the original DNA fragments. These post-ligation fragment sizes reflect an original fragment length between 80 bp–380 bp. Samples underwent sequencing (Princess Margaret Genomics Centre, Toronto, ON, Canada) using the 300-cycle MiSeq Reagent Kit V2 on a MiSeq 2.0 Nano flowcell (Illumina, San Diego, CA, USA), paired-end 2×150 bp, 1 million reads per flowcell (Figure 1).

**Figure 1:**
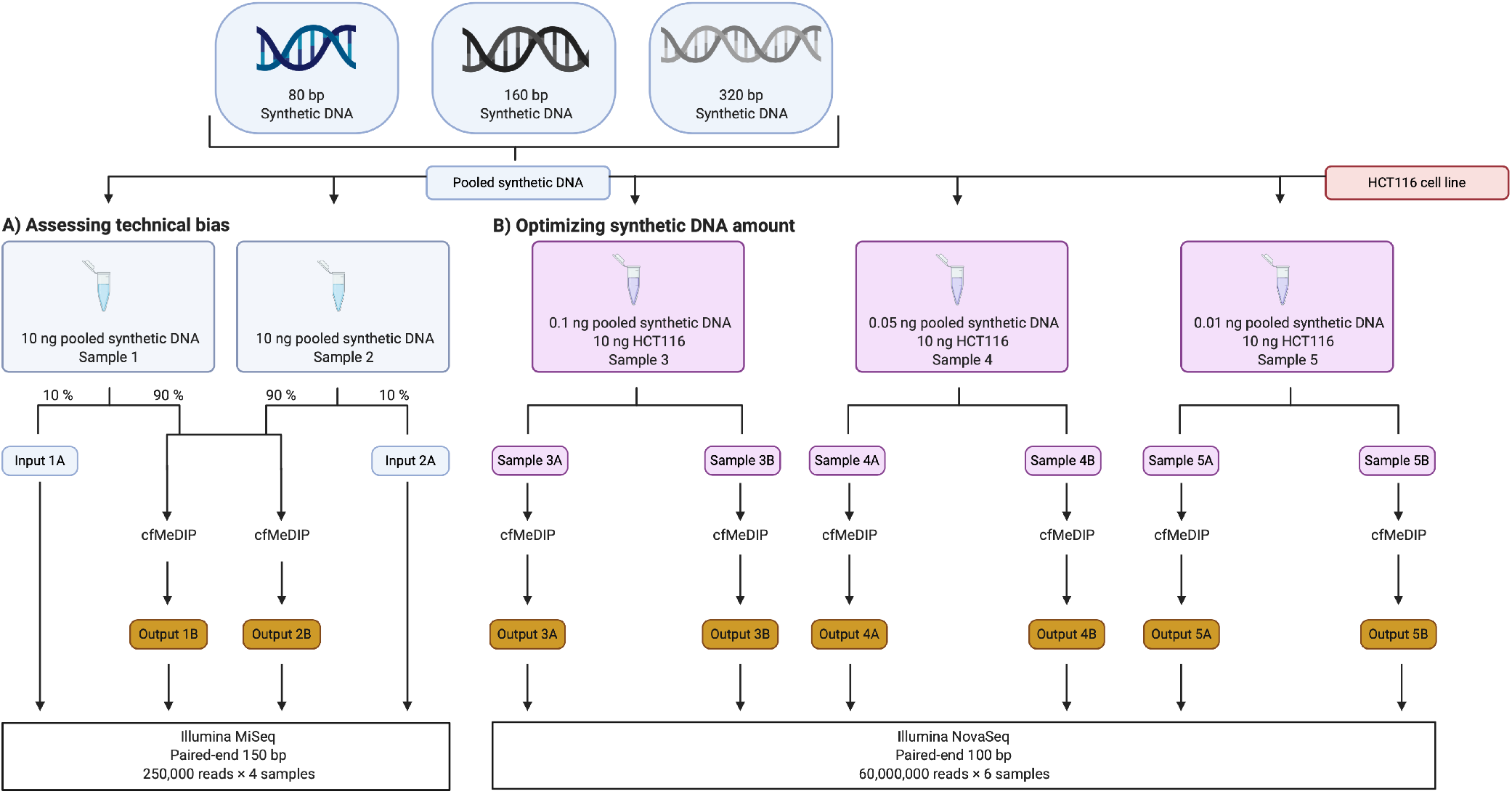
Experimental design using synthetic spike-in control DNA to (A) assess technical bias and (B) optimize the synthetic DNA amount.

### 2.4 Optimizing synthetic DNA amount

We determined the optimal amount of spike-in control DNA needed per experiment by adding varying amounts of spike-in controls to sheared mycoplasma-free HCT116 genomic DNA (American Type Culture Collection, Manassas, VA, USA, RRID: CVCL_0291). Optimizing the amount of spike-in control DNA added to an experiment avoids using a large portion of the sequencing reads on spike-in fragments, saving most reads for the biological sample. We sheared the HCT116 genomic DNA using an LE220 ultrasonicator (Covaris, Woburn, MA, USA). Using AMPure XP beads (Beckman Coulter, Brea, CA, USA), we size selected to ∼ 150 bp in length to mimic cfDNA input. We created 3 replicate samples of sheared HCT116 cfDNA mimic with masses of synthetic spike-in control DNA of 0.1 ng, 0.05 ng, and 0.01 ng. We performed the cfMeDIP-seq experiment as previously described.^2^ Samples underwent sequencing (Princess Margaret Genomics Centre, Toronto, ON, Canada) on a NovaSeq 6000 (Illumina, San Diego, CA, USA), paired-end 2×100 bp, 60 million paired-end reads per sample (Figure 1).

### 2.5 Bioinformatic preprocessing

We performed the same bioinformatic preprocessing on all samples from all experiments. We trimmed adapters using fastp version 0.11.5^17^ `--umi --umi_loc=per_read --umi_len=5 --adapter_sequence=AATGATACGGCGACCAC CGAGATCTACACATATGCGCACACTCTTTCCCTACACGAC --adapter_sequence_r2=CAAGCAGAAGACGGCATACGAGATACGATCAGGTGACTGGAGTTCAGACGTGT`. We removed reads with a Phred score^18^ <40, the default of Bowtie2, and reads with mapping quality <20. We aligned reads to the sequences of our designed fragments using Bowtie2^19^ version 2.3.5 `bowtie2 --local --minins 80 --maxins 320`, writing unaligned reads to a separate file. We subsequently aligned the previous unaligned reads to our synthetic DNA to the human reference genome (GRCh38/hg38).^14^ In every sample, over 98% of reads aligned to either spike-in control sequences or to the human genome. We discarded read pairs when at least one read in the pair did not align or had low quality, defined as Phred score^18^<40 or a mapping quality score <20. We collapsed fragments with the same UMI files counting them as one fragment using UMI-tools version 1.0.0.^20^ We used samtools^21^ version 0.10.2 to convert sequence alignment/map (SAM) files to binary alignment/map (BAM) files.^21^

### 2.6 Absolute quantification from spike-in control data

We created a Gaussian generalized linear model to predict molar amount from deduplicated spike-in control read counts based on UMI consensus sequence, G+C content, CpG fraction, and fragment length. Using this method, we can absolutely quantify cfDNA. To do this, we used the stats package in R^22^ version 3.4.1. Our model estimates molar amount *η* for each DNA fragment present in the original sample using regression coefficients *β* learned for each experiment. For each fragment, this model directly includes read count *x*_reads_, fragment length *x*_len_, G+C content *x*_GC_, and CpG fraction *x*_CpG fraction_. The final model estimates the molar amount *η*,

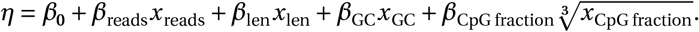

To reduce the left skew of CpG fraction, we used a cube root transformation.

To calculate the proportion of a given fragment that overlapped with the defined 300 bp windows, we used bedtools version 2.29.2 intersect.^23^ We calculated an adjusted molar amount *η*^′^ to only consider the portion of the window each fragment overlapped. For this calculation, we multiplied the molar amount *η* by the length of the overlap between the fragment and the genomic window *ℓ* ∈ [1 bp, 300 bp], divided by the window size *ℓ*^*^ = 300 bp:

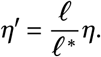

### 2.7 Identifying regions to remove during filtering

To assess multimapping reads that might influence the molar amount estimate, we used Umap^24^ multi-read mappability scores. We used k100 mappability scores, representing the largest read lengths available. We annotated each 300 bp window with its minimum mappability score. We assessed the relationship between molar amount and mappability scores.

We calculated standard deviation of molar amount between the two replicate samples for which we spiked 0.01 ng of synthetic DNA into 10 ng of HCT116 genomic DNA. We assessed the relationship between standard deviation of molar amount between replicates, excluding 1 070 387 simple repeat regions,^25^ 239 461 regions listed in the Encyclopedia of DNA Elements (ENCODE) Project^26^ blacklist,^27^549 876 regions with mappability score ≤0.5, and 906 regions with standard deviation ≥0.05. This left 4 446 375 genomic windows in the analyses.

### 2.8 Correlation between picomoles and M-values

We removed simple repeat regions,^25^ regions listed in the ENCODE blacklist,^27^ regions with Umap k100-multi-read mappability ≤0.5, and regions with standard deviation of molar amount ≥0.05. As described above, we estimated molar amount, using a generalized linear model (*r*^2^ = 0.93). We prioritized models that performed better on 160 bp fragments, as we physically size selected for these fragments.

To compare molar amount to another complimentary measure of DNA methylation, we had genomic DNA from the cell line HCT116 profiled (Princess Margaret Genomics Centre, Toronto, ON, Canada) using the Infinium MethylationEPIC BeadChip array (Illumina, San Diego, CA, USA). We prepared these samples as technical replicates of the HCT116 genomic DNA that we later spiked with 0.01 ng spike-in control. We normalized and preprocessed array data using sesame^28^ version 1.8.2. We annotated CpGs on the array to our 300 bp genomic windows. When >1 CpG probe annotated to a window, we calculated the mean probe M-values across the window.

We assessed correlation between array M-values and picomoles, and between array M-values and read counts. We compared to M-values rather than *β* values because *β* values have high heteroscedasticity.^29^ As cfMeDIP-seq preferentially enriches for highly methylated regions, we hypothesized that regions for which the array has more CpG probes would correlate to both molar amount and read counts better than regions with less CpG coverage. As such, we assessed the correlation independently at windows containing ≥3, ≥5, ≥7, and ≥10 CpG probes.

### 2.9 Examining consistency across experimental batches

To experimentally introduce batch effects, we provided a sample of 10 ng of cfDNA obtained from peripheral blood plasma of 5 acute myeloid leukemia (AML) patients containing 0.01 ng of our spike-in controls to 3 independent labs. We obtained the deidentified patient samples, previously included in Shen *et al*.,^1^ from the Leukemia Tissue Bank, Princess Margaret Cancer Centre, University Health Network with informed consent following approval by the University Health Network Research Ethics Board (01-0573). Blood was collected at the time of diagnosis in EDTA tubes. Samples were spun and plasma frozen in Eppendorf tubes at −80 °C until use. As a blood cancer, leukemia generates a high amount of plasma cfDNA, allowing us to have sufficient cfDNA to divide into three technical replicates of 10 ng each.

Each lab performed the cfMeDIP-seq method described by Shen et al,^2^ with a variety of procedural modifications, detailed here. The changes emulated batch effects commonly found in publicly available data from different labs. Labs 1 and 3 used the same IDT xGen Duplex Seq adapters with 3 bp–4 bp UMI, as described above. Lab 2 used custom IDT adapters with 2 bp degenerate UMI, as previously described.^30^ For ligation of adapters, Labs 1 and 2 incubated at 4 °C for 16 h, while Lab 3 incubated at 20 °C for 2 h. Labs 1 and 3 used Antibody 1 (Diagenode, Denville, NJ, USA, Cat #C15200081-100, Lot #RD004, RRID: AB_2572207), while Lab 2 used Antibody 2 (Diagenode, Denville, NJ, USA, Cat #C15200081-100, Lot #RD001, RRID: AB_2572207). Lab 1 and 2 used 50% methylated lambda filler DNA and 50% unmethylated lambda filler DNA. Lab 3 used 100% unmethylated lambda filler DNA.

For amplifying the final library, the batches had different numbers of PCR cycles. Lab 1 ran 13–15 cycles, optimized per sample, Lab 2 ran 13 cycles, and Lab 3 ran 11 cycles. Lab 1 and 2 sequenced DNA with a NovaSeq 6000 (Illumina,

San Diego, CA, USA), paired-end 2×100 bp. Lab 3 sequenced DNA on a NextSeq 550 (Illumina, San Diego, CA, USA), paired-end 2×75 bp. Lab 1 obtained 60 million reads per sample, Lab 2 obtained 100 million reads per sample, and Lab 3 obtained 85 million reads per sample.

We assessed whether spike-in controls mitigated batch effects on samples for which we calculated molar amount. To do this, we performed principal component analysis (PCA) on four different quantification methods:

1. read counts only, without the use of spike-in controls
2. read counts preprocessed using QSEA version 1.16.0,^31^ the current standard processing pipeline for the methylated DNA immunoprecipitation-sequencing (MeDIP-seq) data
3. molar amount, without filtering
4. molar amount, with filtering for repeat regions, ENCODE blacklist, and regions with low mappability

To investigate whether known variables associated with any of the principal components, we performed two-way analysis of variance (ANOVA) between each principal component and each categorical variable. Categorical variables included batch, filler (methylated or unmethylated), adapter, samples, and sex (inferred by Y chromosome signal). We converted the resulting F-statistic to an effect size, Cohen’s *d*,^32^ using the compute.es package^33^ version 0.2.5 and R version 3.4.1. We adjusted p-values for multiple test correction using the Holm-Bonferroni method.^34^

## 3 Results

### 3.1 Spike-in controls confirm cfMeDIP’s high efficiency and specificity

We performed cfMeDIP-seq directly on the synthetic spike-in control fragments. For each sample, we saved 10% of the mass before performing cfMeDIP-seq. This acts as an input control (Figure 1). In input samples, we observed 51% of the input fragments methylated and 49% unmethylated. After cfMeDIP, 97% of the output fragments were methylated (Figure 2A,B). The enrichment for methylated sequences further supports the validity and high efficiency of cfMeDIP-seq.

**Figure 2:**
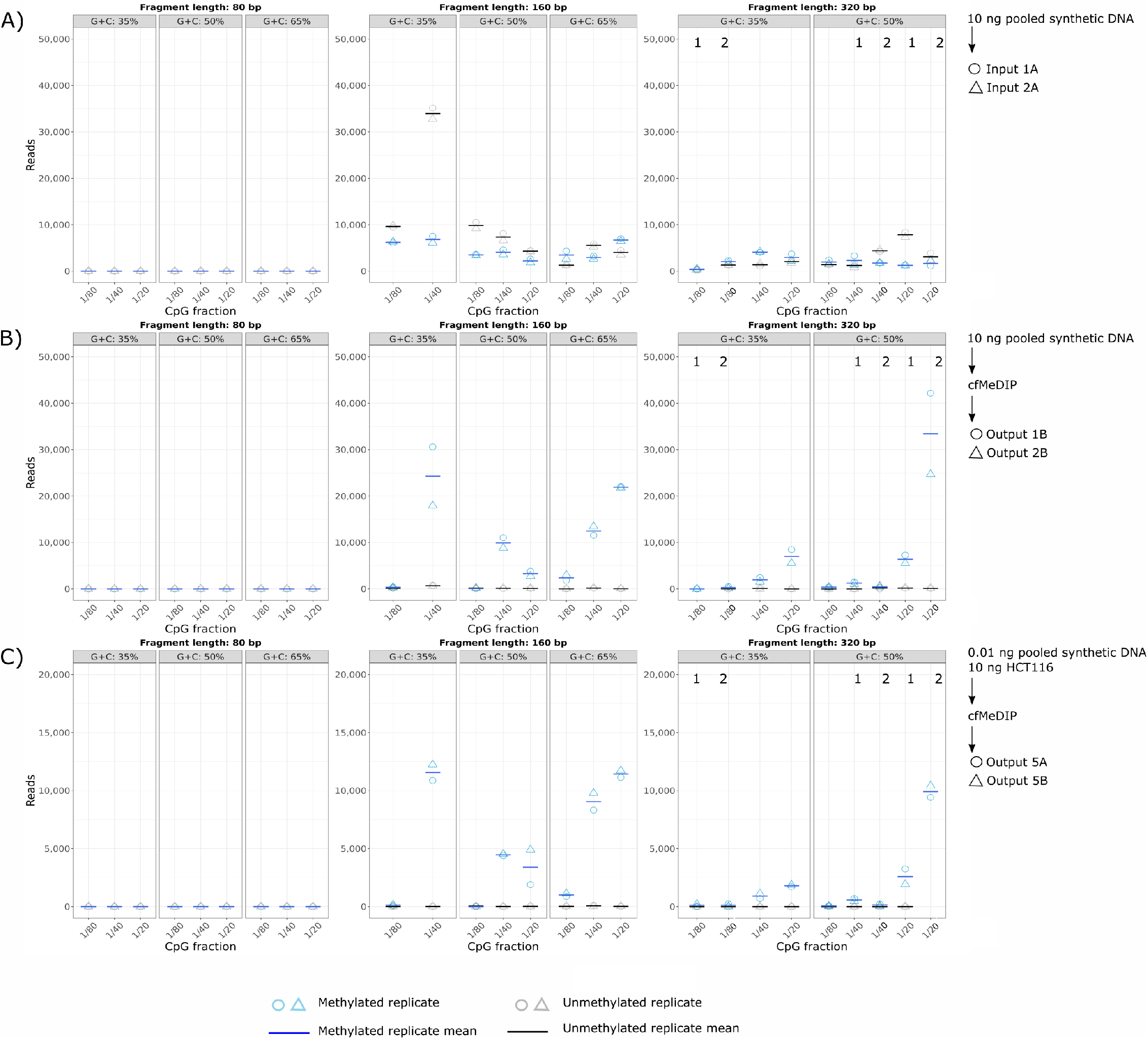
Assessing biases in fragment length, G+C content, and CpG fraction in (A) input spike-in control DNA without cfMeDIP-seq, (B) output spike-in control DNA, after cfMeDIP-seq, and (C) 0.01 ng spike-in control DNA added to HCT116 replicates. Blue: methylated fragments; grey: unmethylated fragments. Circle: sample 1; triangle: sample 2. Solid line: mean of the two samples. Columns marked with numerals 1 and 2 represent alternative sets of fragments with identical properties but different sequences.

After cfMeDIP, signal from methylated fragments for both the synthetic spike-in control alone (97% of spike-in control fragments) and 10 ng HCT116 DNA with 0.01 ng spike-in (97% of spike-in control fragments) showed an enrichment of 160 bp fragments. We expected this enrichment due to our size selection step for insert size fragments between 80 bp–380 bp. We also observed enrichment of fragments with higher G+C content and higher CpG fraction (Figure 2B,C).

Signal from unmethylated fragments for both the synthetic spike-in control alone (3% of spike-in control fragments) and 10 ng HCT116 DNA with 0.01 ng spike-in (3% of spike-in control fragments) had no association with fragment lengths, G+C content, or CpG fraction (Figure 2B,C). This suggests that the low amount of unmethylated fragment signal arose from random non-specific binding and that it was not confounded systematically with fragment length, G+C content, or CpG fraction.

We assessed the range of number of sequenced reads where we had alternative sets of 320 bp fragments with different G+C contents and CpG fractions. Because no unmethylated fragment yielded more than 20 reads in any experiment, we focused our assessment on methylated fragments. The similar numbers of reads sequenced for alternate fragments with the same fragment properties show the robustness of the spike-in controls (Table 1).

**Table 1:** Range of reads sequenced for alternative synthetic spike-in fragments length 320 bp with the same G+C contents and CpG fractions. ^a^Minimum and maximum range between the individual replicates of two alternatives. ^b^Range between the means of two alternatives.

| Experiment | G+C content | CpG fraction | Within-alternative<sup>a</sup> | Between alternatives<sup>b</sup> |
| --- | --- | --- | --- | --- |
| 10 ng synthetic DNA | 35% | 1/80 | 0–343 reads | 343 reads |
|  | 50% | 1/40 | 298–445 reads | 1161 reads |
|  | 50% | 1/20 | 1759–17 472 reads | 36642 reads |
| HCT116 + 0.01 ng synthetic DNA | 35% | 1/80 | 13–219 reads | 219 reads |
|  | 50% | 1/40 | 160–240 reads | 646 reads |
|  | 50% | 1/20 | 987–1320 reads | 8483 reads |

Even in the fragments with the largest range between alternatives (50% G+C content, 1/20 bp CpG fraction), this range represents only ≤ 4% of the total number of sequenced reads per sample. This difference may represent unknown fragment properties that influences cfMeDIP-seq results. The other sets of alternative fragments had even smaller ranges between alternatives.

### 3.2 Low-input spike-in control accounts for technical variance

To determine the optimal amount of spike-in control DNA to add to each sample, we assessed the the proportion of reads used towards our spike-in controls. We compared spike-in control reads to the total number of reads used for our biological sample, HCT116 genomic DNA. We optimized the amount of spike-in controls to be used in subsequent experiments. This allowed us to maximize reads to our biological sample of interest while obtaining sufficient reads from the spike-in controls to correct for technical bias. Adding in 0.01 ng of our synthetic spike-in control DNA before cfMeDIP-seq used <1% of the reads for the spike-in controls. We retained >650 000 reads of spike-in control sequence for analysis while leaving the rest of the reads for our biological sample. Therefore, we decided to use 0.01 ng of spike-in control fragments in subsequent experiments.

The 0.01 ng of spike-in control DNA added to 10 ng of HCT116 genomic DNA revealed ≥97% specificity of methylated

DNA with ≤ 0.01% total non-specific binding to non-methylated fragments (Figure 2C). We calculated methylation specificity by dividing the total number of methylated fragments by the total number of spike-in control fragments. The cfMeDIP process also enriched for the 160 bp fragments we physically size selected for and for higher G+C content. Fragments that have CpG present at only1/80 bp in the experiment with 10 ng of HCT116 with 0.01 ng of spike-in control DNA were represented by ≤ 1% of reads (Figure 2C). The same patterns persisted when spiking in 0.05 ng and 0.1 ng spike-in control DNA into 10 ng of HCT116 cell line.

### 3.3 Removing problematic regions eliminates some technical artifacts

From our normalized data, we removed regions containing simple repeats, the ENCODE blacklist regions, and regions with mappability scores ≤0.5. We noticed that many regions with high molar amount also had high standard deviation of molar amount between replicate samples. Thus, we removed regions where standard deviation of molar amount between replicates was ≥0.05.

After removing the high standard deviation regions, we observed no relationship between molar amount and standard deviation of molar amount, and no relationship between molar amount and mappability (Figure 3). We further examined the 11 genomic windows with the highest estimated molar amounts (≥ 0.25 pmol; Figure 3A,B). Of these 11 windows, 7 overlapped repetitive elements (Table 2). Of those 7, 6 overlapped with Alu elements. This suggests that removing regions that have high standard deviation of molar amount between replicates can reduce technical artifacts. Regions with high standard deviation of molar amount between replicates perform inconsistently, leading to inaccurate quantification of DNA methylation.

**Figure 3:**
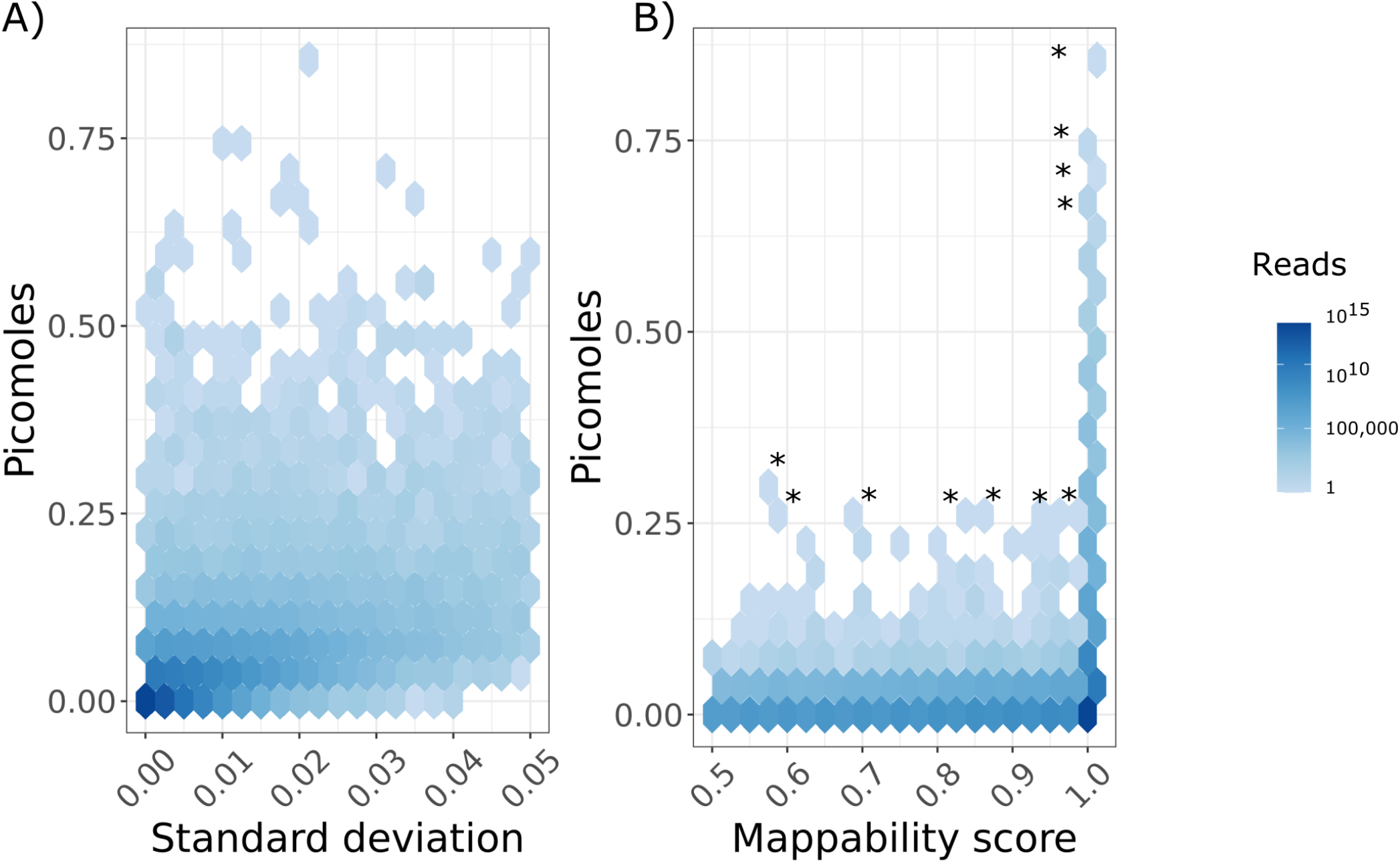
Two-dimensional histogram of the number of reads found in 300 bp windows, as binned by molar amount and either (A) standard deviation of molar amount or (B) Umap k100 multi-read mappability. Only includes windows that do not overlap with University of California, Santa Cruz (UCSC) simple repeats and the ENCODE blacklist, and regions with Umap k100 multi-read mappability scores ≤0.5. * 11 outlier genomic windows chosen for further examination.

**Table 2:**
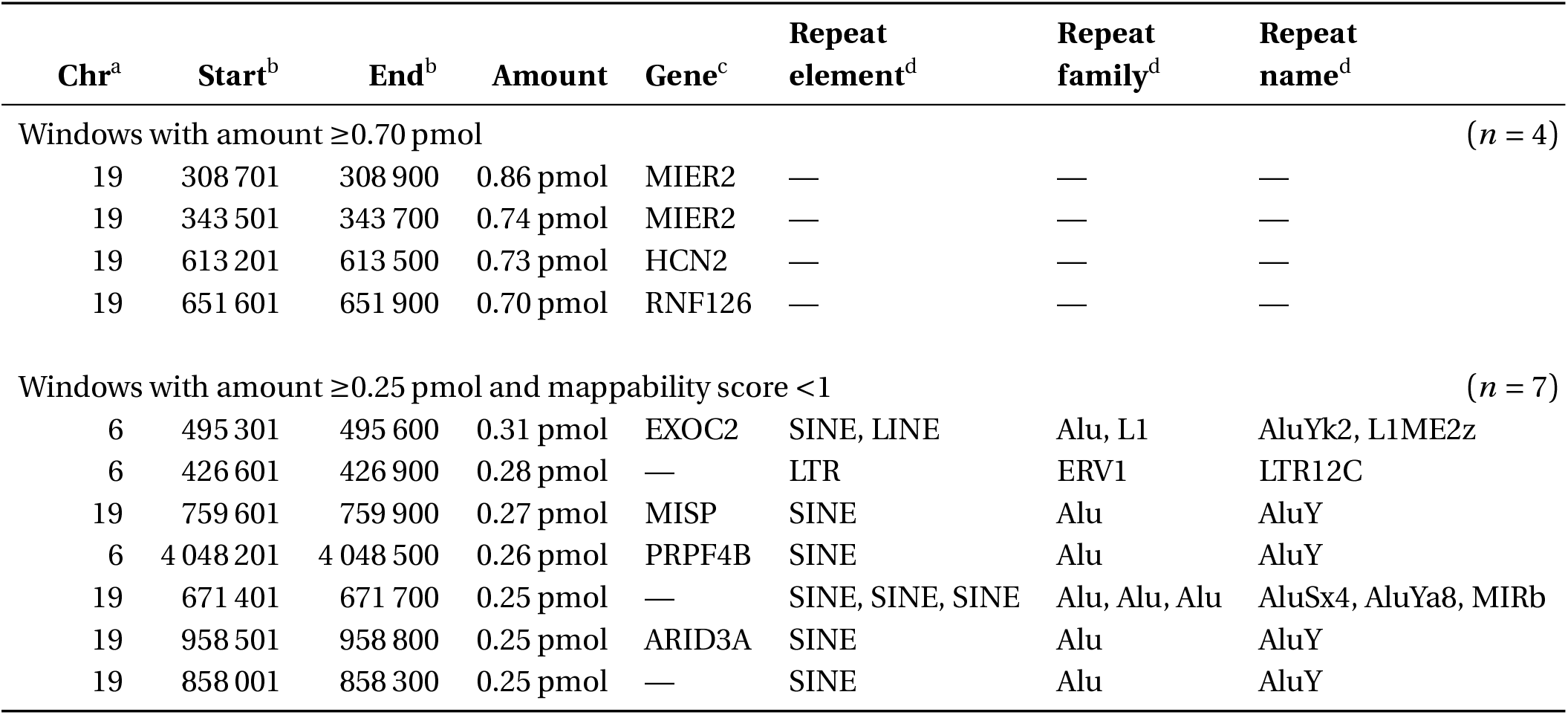
11 genomic windows of length 300 bp with high predicted molar amount. Sorted by decreasing molar amount. ^a^Chromosome. ^b^GRCh38/hg38, genomic position 1-start, fully closed. ^c^Symbols of GENCODE version 33 basic gene set genes35 that overlap our 300 bp genomic windows. ^d^Elements, families, and names of RepeatMasker36 version 3.0 repeats that overlap our 300 bp genomic windows.

| Chr<sup>a</sup> | Start<sup>b</sup> | End<sup>b</sup> | Amount | Gene<sup>c</sup> | Repeat element<sup>d</sup> | Repeat family<sup>d</sup> | Repeat name<sup>d</sup> |
| --- | --- | --- | --- | --- | --- | --- | --- |
| Windows with amount $\geq 0.70$ pmol | | | | | | | |
| 19 | 308 701 | 308 900 | 0.86 pmol | MIER2 | — | — | — |
| 19 | 343 501 | 343 700 | 0.74 pmol | MIER2 | — | — | — |
| 19 | 613 201 | 613 500 | 0.73 pmol | HCN2 | — | — | — |
| 19 | 651 601 | 651 900 | 0.70 pmol | RNF126 | — | — | — |
| Windows with amount $\geq 0.25$ pmol and mappability score $< 1$ | | | | | | | |
| 6 | 495 301 | 495 600 | 0.31 pmol | EXOC2 | SINE, LINE | Alu, L1 | AluYk2, L1ME2z |
| 6 | 426 601 | 426 900 | 0.28 pmol | — | LTR | ERV1 | LTR12C |
| 19 | 759 601 | 759 900 | 0.27 pmol | MISP | SINE | Alu | AluY |
| 6 | 4 048 201 | 4 048 500 | 0.26 pmol | PRPF4B | SINE | Alu | AluY |
| 19 | 671 401 | 671 700 | 0.25 pmol | — | SINE, SINE, SINE | Alu, Alu, Alu | AluSx4, AluYa8, MIRb |
| 19 | 958 501 | 958 800 | 0.25 pmol | ARID3A | SINE | Alu | AluY |
| 19 | 858 001 | 858 300 | 0.25 pmol | — | SINE | Alu | AluY |

### 3.4 Absolute quantification correlates with M-values

We compared molar amount to M-values from the EPIC array. We used our generalized linear model to calculate molar amount of cfMeDIP-seq methylated DNA fragments for each 300 bp genomic window. Molar amount significantly correlated with array M-values over 300 bp in our HCT116 genomic DNA samples (*r* ≥ 0.62; Figure 4A,C,E,G). Correlation increased when we restricted analyses to 300 bp windows with ≥5 CpG probes on the array (*r* = 0.82 ; Figure 4C). This reflected cfMeDIP-seq’s preference for methylated, CpG-dense reads.

**Figure 4:**
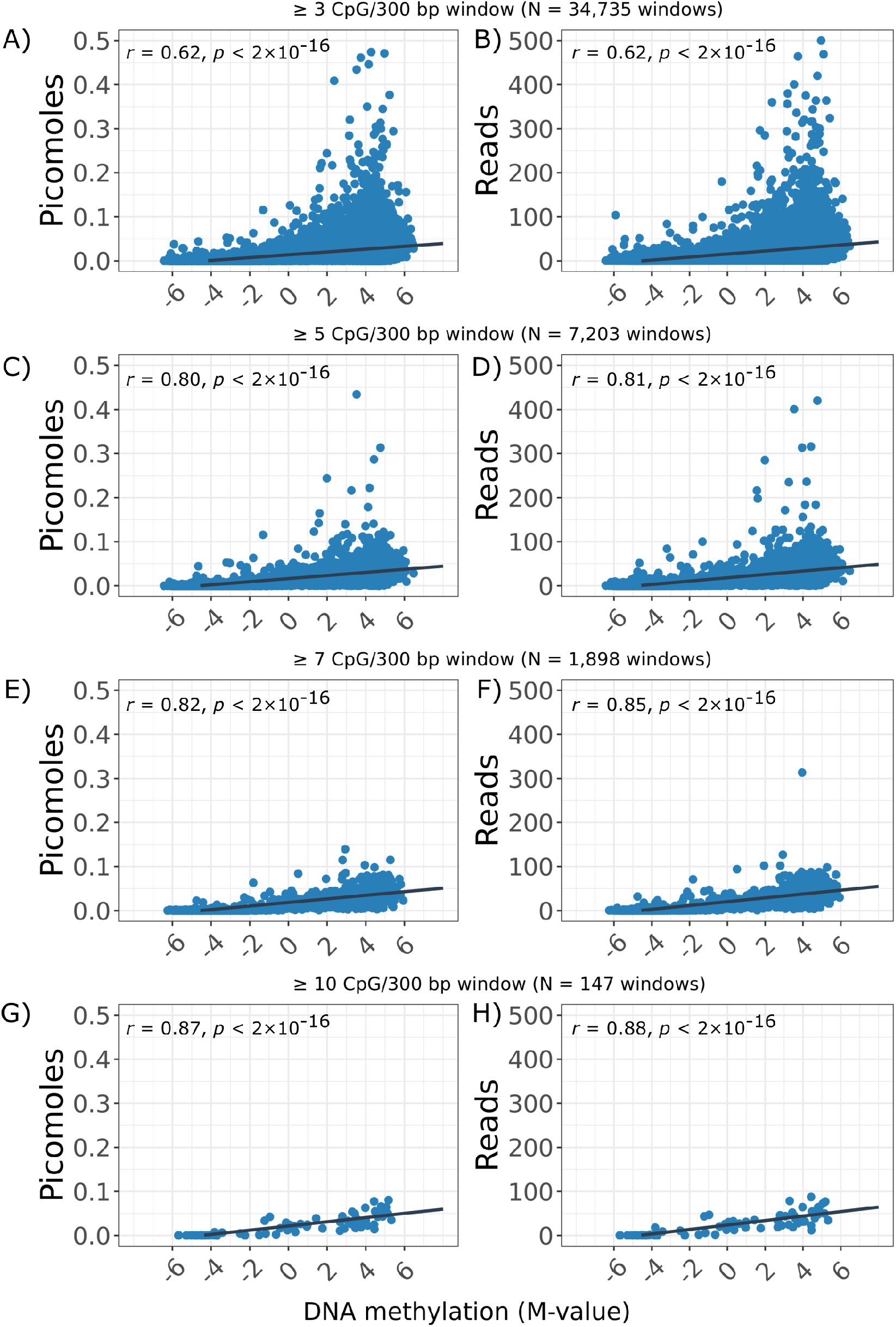
Correlation of two measurements of fragment methylation by cfMeDIP and EPIC array M-value for 300 bp genomic windows. (A,C,E,G) Molar amount calculated from HCT116 samples correlated to EPIC array M-values. (B,D,F,H) Read counts calculated from the same samples, ignoring the spike-in controls. (A,B) 37 714 windows with ≥3 CpG probes represented on the EPIC array. (C,D) 7975 windows with ≥5 CpG probes represented on the EPIC array. (E,F) 2066 windows with ≥7 CpG probes represented on the EPIC array. (G,H) 158 windows with ≥10 CpG probes represented on the EPIC array.

We compared the current standard of read counts to M-values (Figure 4B,D,F,H). Molar amount correlated with

M-values similarly to read counts, but provides the advantage of absolute quantification.

### 3.5 Spike-in controls significantly mitigate batch effects

To determine whether our spike-in controls mitigate batch effects, we provided aliquots of cfDNA samples from 5 AML patients to three different labs. Each lab performed cfMeDIP-seq on each of the 5 samples with slight variations.

We performed PCA to assess if any batch variables drive any of the top principal components. For raw read count data without spike-in controls, principal component 1 explains 80% of the variance and showed DNA methylation changes non-significantly associated with processing batch, filler (methylated or unmethylated), and adapter. For the raw reads, principal component 3, explaining 5% of the variance, significantly associated with batch (Figure 5A).

**Figure 5:**
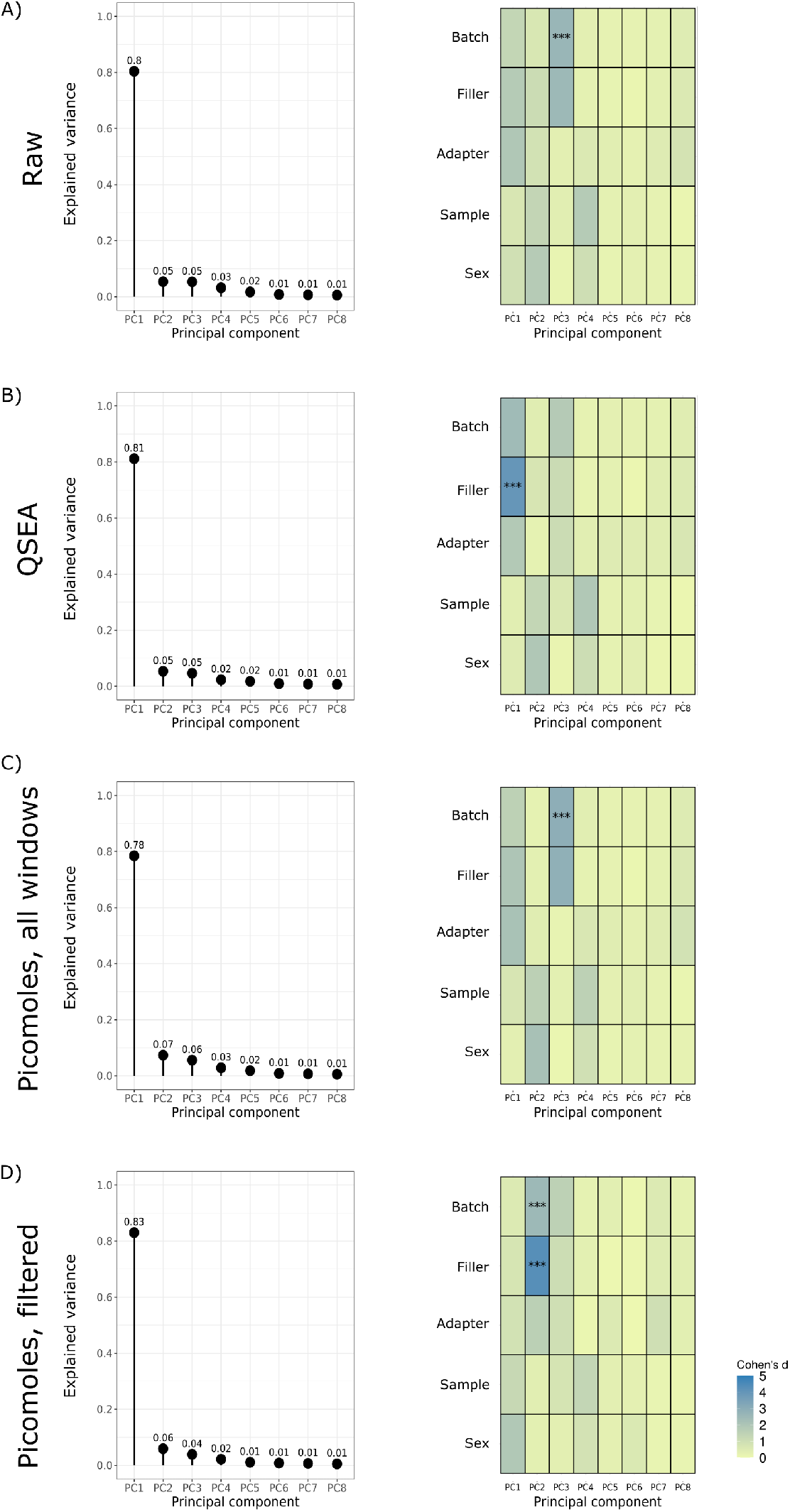
Principal component analyses of cfMeDIP results normalized through 4 different strategies, and as-sociations with experimental variables. *(Left)* Proportion of the variance explained by each principal component. *(Right)* Association between known variables, both technical and clinical, and principal component. Cohen’s *d* is an effect size of standardized means between variable. *** *p* < 0.001. **(A) Raw read counts without any normalization. (B) Read counts normalized using QSEA. (C) Data normalized using spike-in controls. (D) Data normalized using spike-in controls and removing regions in UCSC simple repeats, in the ENCODE blacklist, and with Umap k100 multi-read mappability scores** ≤**0.5**.

QSEA normalization increased the variance explained by principal component 1 to 81%. With QSEA normalization, principal component 1 significantly associated with filler. It also associated non-significantly with batch and adapter (Figure 5B). The correlation of principal component 1 to filler DNA only appeared after QSEA normalization, suggesting that QSEA may have introduced stochastic variance into the data.

Using spike-in controls greatly reduced the variance explained by principal component 1 to 78%. Similar to the raw data, we still see association with the batch variable in principal component 3, explaining 6% of the variance (Figure 5C).

Using our suggested filtering, which includes removing simple repeats, ENCODE blacklist regions, and regions with low mappability, improves variance associated with batch variables in the data. Upon removing these regions, batch and filler associated with principal component 2, explaining 6% of the variance. These variables were not confounded with the known non-technical, biological variables—sample and sex—that one would want to associate with the top principal components. Principal component 1, explaining 83% of the variance, associated with sample and sex more than other variables, though not significantly (Figure 5D). These changes in DNA methylation associated with biological variable were absent in principal component 1 of either the raw data (Figure 5A) or the QSEA data (Figure 5B).

We further investigated principal component 2 using spike-in controls and filtering. Examining the top 10% of windows (*N* = 258254) driving the explained variance, 71% matched to RepeatMasker^36^ repetitive elements. Of the mapped repetitive elements, 43% mapped to Alu elements. Batch effects associated with repetitive elements suggest inconsistent or inaccurate measures of DNA methylation at these regions.

## 4 Discussion

The data above establish the validity of using our synthetic spike-in control DNA to absolutely quantify cfDNA in cfMeDIP-seq experiments. We showed that technical bias exists in the cfMeDIP-seq data, and that the use of our spike-in controls helps mitigate these biases. In cfMeDIP-seq data generated without using the spike-in controls, the batch effects may prove stark. For example, Lab 3 used unmethylated lambda filler DNA, but the spike-in controls successfully mitigated the effect of this change. One can use the signal from unmethylated spike-in control reads to calculate methylation specificity for each sample. To reduce technical artifacts, one must remove problematic genomic regions prior to analysis.

Despite removing problematic genomic regions, some remaining regions had much higher predicted molar amount than all other genomic regions of a given standard deviation or mappability score. The high molar amount genomic regions consisted mostly of repetitive elements, predominantly short interspersed nuclear elements (SINEs) (Table 2). While these regions had high CpG density, our model already adjusted for CpG fraction. This made CpG density an unlikely driver of high molar amount.

The two highest molar amount genomic regions from our HCT116 colon cancer cells overlap *MIER2*, part of the MIER gene family, associated with suppression of colorectal cancer.^37^ This shows the utility of our method to reveal important pathways for the gene regulation of the sample examined by a liquid biopsy.

Out of the top 5 genomic regions we followed up, 4 had high expression in the testis (*MIER2, RNF126, EXOC2*), and 1 had high expression in the ovary (*PRPF4B*). Several cancers exhibit high expression of genes associated with cell lines and low methylation in these genes’ promoters.^38–40^

We may see over-representation of high molar amount in some repetitive regions due to the characteristic hypermethylation of these regions.^41^ Had we not removed many repetitive regions when removing regions listed in the ENCODE blacklist and regions with low mappability, we may have observed more genomic regions with predicted high molar amount. Regions with high molar amount that map to repetitive elements contain many extra copies not present in the reference genome. These regions would appear uniquely mappable, and thus not removed by our previous filtering steps.

HCT116 likely has problematic genomic regions not found in the ENCODE blacklist. This arises from the relative dearth of ENCODE data available for the blacklist generation process, compared to cell types in the ENCODE Tier 1 and Tier 2 categories.

Depending on the experimental question, some may choose to go beyond our filtering recommendations and remove all repetitive elements, such as all long interspersed nuclear elements (LINEs) and all SINEs. Given that, after filtering, only 11 genomic windows had molar amount ≥ 0.25 pmol, removing all repetitive elements would not affect results drastically.

Both molar amount in picomoles and read counts correlated to M-values. The strength of this correlation varied with the number of CpGs represented on the EPIC array for a genomic window. We expect the increase in correlation with more CpGs as cfMeDIP-seq preferentially enriches for hypermethylated regions, and picks up fewer fragments in CpG-sparse regions. Additionally, if the array has fewer probes representing a 300 bp genomic region than that region has CpGs, it may poorly represent DNA methylation for the entire 300 bp.

To facilitate the use of our spike-in controls, we have created an R package, `spiky`, to help process data generated from cfMeDIP-seq experiments that include the spike-in controls. This package trains the Gaussian generalized linear model and predicts molar amount in picomoles on user data. The `spiky` package is available on GitHub (https://github.com/trichelab/spiky) and will soon be available on Bioconductor.^42^

Incorporating these spike-in controls in future cfMeDIP-seq experiments will adjust for technical biases and mitigate batch effects, improving cfMeDIP-seq data overall.

## Data availability

We deposited processed data and raw non-human data in the Gene Expression Omnibus^43^ (GEO accession: GSE166259). We deposited raw data for the AML samples in the European Genome-phenome Archive^44^ (EGA accession: EGAS00001005069), with a specified policy for processing data access requests (https://doi.org/10.5281/zenodo.4568265).

## Code availability

The `spiky` package is available on GitHub (https://github.com/trichelab/spiky) under the GNU General Public License version 2 license. Other scripts are available on GitHub (https://github.com/hoffmangroup/2020spikein) and deposited on Zenodo (https://doi.org/10.5281/zenodo.4533340).

## Author contributions

Conceptualization, S.L.W., S.Y.S., D.D.De C., and M.M.H.; Data curation, S.L.W. and S.Y.S.; Formal analysis, S.L.W.; Funding acquisition, S.L.W., S.V.B., D.D.De C., and M.M.H.; Investigation, S.Y.S. and J.B.; Methodology, S.L.W., S.Y.S., T.T., D.D.De C., and M.M.H.; Project administration, S.L.W. and M.M.H.; Resources, M.M.H.; Software, S.L.W., L.H., and T.T.; Supervision, S.V.B., T.T., D.D.De C., and M.M.H.; Validation, S.L.W. and S.Y.S.; Visualization, S.L.W.; Writing — original draft, S.L.W.; Writing — review & editing, S.L.W., S.Y.S, L.H., J.B., T.T., S.V.B., D.D.De C., and M.M.H.

## Acknowledgments

We thank Dax Torti (0000-0002-9000-5803) (Ontario Institute for Cancer Research) for performing batch analysis experiments, Mark D. Minden (0000-0002-9089-8816) and Andrea Arruda (0000-0003-2516-8005) (Leukemia Tissue Bank, Princess Margaret Cancer Centre, University Health Network) for providing the AML samples and for comments on the manuscript, and Zhibin Lu (0000-0001-6281-1413) (Bioinformatics and High Performance Computing Core, University Health Network) for technical assistance. Figure 1 created with BioRender.com, which has a restrictive license that requires this acknowledgment. This work was supported by the Canadian Institutes of Health Research (389866 to M.M.H., MFE-171256 to S.L.W.), the McLaughlin Centre (MC-2019-07 to M.M.H.), the Molly Towell Perinatal Research Foundation, and the Princess Margaret Cancer Foundation. T.T. is supported by startup funds from the Van Andel Institute, the Michigan Economic Development Corporation, the Michelle Lunn Hope Foundation, and the Folz Family Fund for Cancer Research.

